# Genetic basis of a spontaneous mutation’s expressivity

**DOI:** 10.1101/2020.04.03.024547

**Authors:** Rachel Schell, Joseph J. Hale, Martin N. Mullis, Takeshi Matsui, Ryan Foree, Ian M. Ehrenreich

## Abstract

Genetic background often influences the phenotypic consequences of mutations, resulting in variable expressivity. How standing genetic variants collectively cause this phenomenon is not fully understood. Here, we comprehensively identify loci in a budding yeast cross that impact the growth of individuals carrying a spontaneous missense mutation in the nuclear-encoded mitochondrial ribosomal gene *MRP20*. Initial results suggested that a single large effect locus influences the mutation’s expressivity, with one allele causing inviability in mutants. However, further experiments revealed this simplicity was an illusion. In fact, many additional loci shape the mutation’s expressivity, collectively leading to a wide spectrum of mutational responses. These results exemplify how complex combinations of alleles can produce a diversity of qualitative and quantitative responses to the same mutation.

## Introduction

Mutations frequently exhibit different effects in genetically distinct individuals (or ‘background effects’) (1–3). For example, not all people with the same disease-causing mutations manifest the associated disorder or exhibit identical symptoms. A commonly observed form of background effect among individuals carrying the same mutation is different degrees of response to that mutation (or ‘variable expressivity’) (4). Variable expressivity can arise due to a myriad of reasons, including genetic interactions (or epistasis) between a mutation and segregating loci (1), dominance (1), stochastic noise (5), the microbiome (6), and the environment (1).

The role of epistasis in expressivity has proven especially difficult to study, in part because natural populations harbor substantial genetic diversity, which can facilitate complex genetic interactions between segregating loci and mutations (7–20). Mapping the loci involved in these interactions is technically challenging. However, controlled laboratory crosses provide a powerful tool for identifying the loci that interact with particular mutations, giving rise to background effects (7, 12, 17–19, 21).

In this paper, we use a series of controlled crosses in the budding yeast *Saccharomyces cerevisiae* to comprehensively characterize the genetic basis of a mutation’s expressivity. We focus on a missense mutation in *MRP20*, an essential nuclear-encoded subunit of the mitochondrial ribosome (22). This mutation occurred by chance in a cross between the reference strain BY4716 (‘BY’) and a clinical isolate 322134S (‘3S’), and was found to show variable expressivity among BYx3S cross progeny. This presented an opportunity to determine how loci segregating in the BYx3S cross individually and collectively influence this mutation’s expressivity.

## Results

### A spontaneous mutation increases phenotypic variance in the BYx3S cross

In the BY/3S diploid progenitor of haploid BYx3S segregants, a spontaneous mutation occurred in a core domain of Mrp20 that is conserved from bacteria to humans (Fig. 1A, fig. S1, table S1) (22, 23). This mutation resulted in an alanine to glutamine substitution at amino acid 105 (*mrp20-105E*) and showed variable expressivity among segregants carrying it. Specifically, segregants with this mutation showed increased phenotypic variance relative to wild type segregants when ethanol was provided as the carbon source, the condition used hereafter (Fig. 1B; Levene’s test, p = 5.9 × 10^−22^). Mutant segregants exhibited levels of growth ranging from inviable to wild type, and fit a bimodal distribution that centered on 10% and 57% growth relative to the haploid BY parent strain (bimodal fit log likelihood = 30; fig. S2).

**Figure 1.**
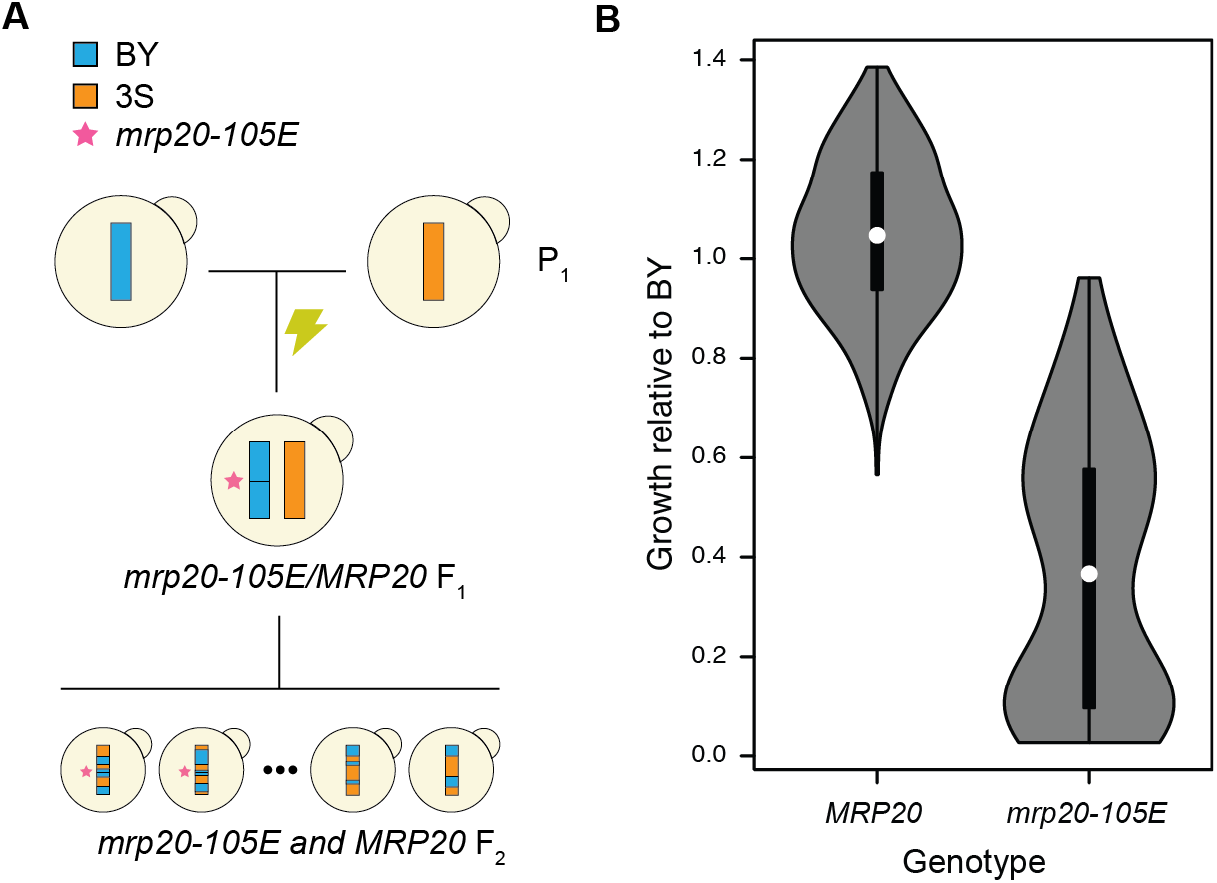
The *mrp20-105E* mutation occurred spontaneously, increasing phenotypic variance in the BYx3S cross. (A) A spontaneous mutation in a BY/3S diploid gave rise to a BYx3S segregant population in which *mrp20-105E* segregated. (B) The *mrp20-105E* segregants exhibited increased phenotypic variance and a bimodal distribution of phenotypes. Throughout the paper, blue and orange are used to denote BY and 3S genetic material, respectively. All growth data presented in the paper are measurements of colonies on agar plates containing rich medium with ethanol as the carbon source.

### A large effect locus shows epistasis with *mrp20-105E*

Loci contributing to this variable expressivity should be detectable through their genetic interactions with *MRP20*. To find such loci, we performed linkage scans for two-way epistasis with *MRP20*. We identified a single locus on Chromosome XIV (ANOVA, interaction term p = 4.3 × 10^−16^; Fig. 2A). Individuals with *XIV^BY^* showed reduced growth among both *MRP20* and *mrp20-105E* segregants, but to a greater degree among the latter (Fig. 2B). The Chromosome XIV locus explained 79% of the phenotypic variance among *mrp20-105E* segregants (ANOVA, p = 3.2 × 10^−31^) and accounted for all observed cases of inviability (Fig. 2B).

**Figure 2.**
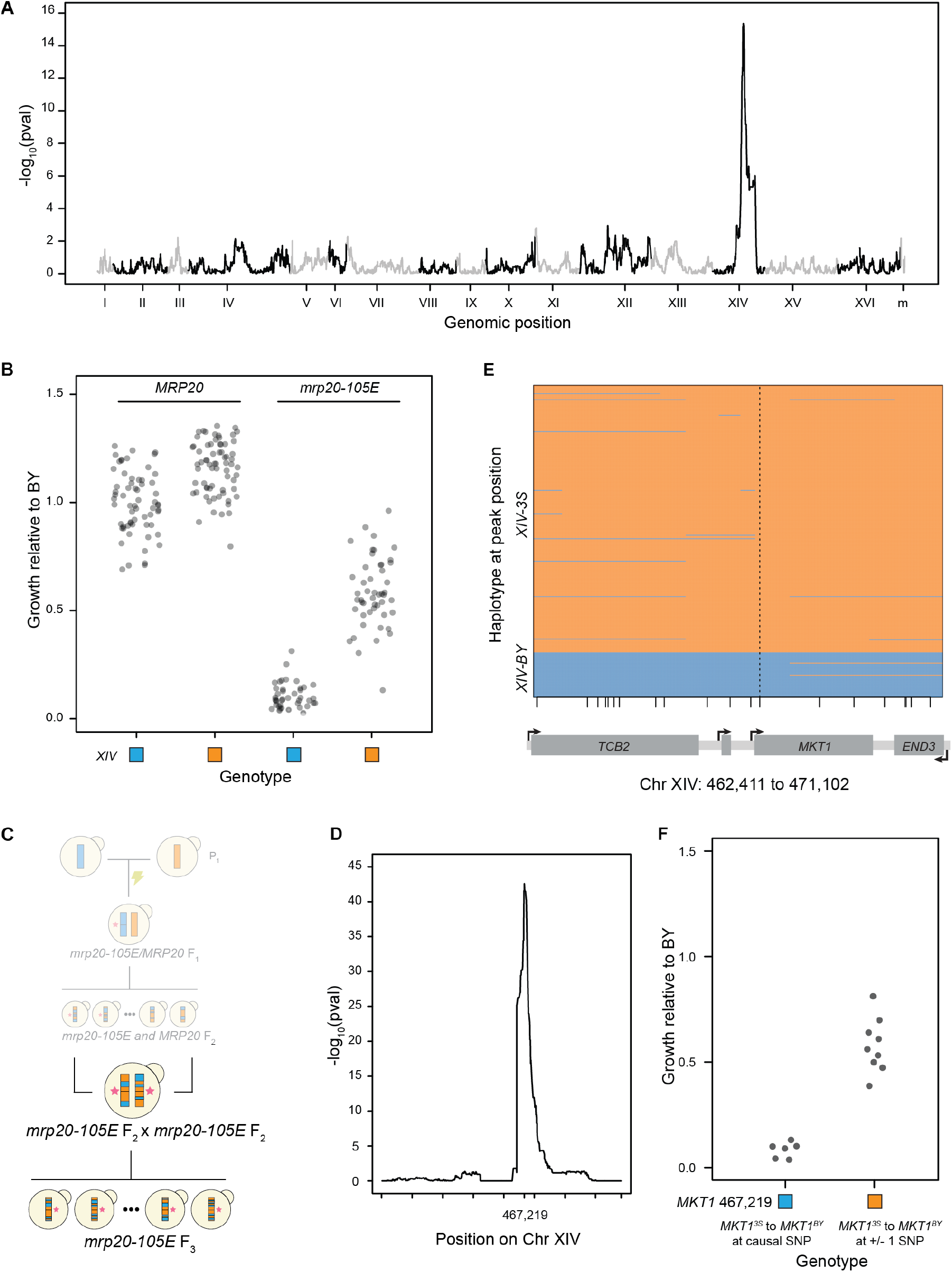
Epistasis between *MRP20* and *MKT1* appears to mostly explain response to *mrp20-105E*. (A) Linkage mapping in the BYx3S segregants shown in Fig 1 identified a locus on Chromosome XIV that exhibits a two-way genetic interaction with *MRP20*. (B) The Chromosome XIV locus had effects in both *MRP20* and *mrp20-105E* segregants but had a greater effect among *mrp20-105E* segregants. (C) To identify the causal gene, we crossed two *mrp20-105E* F_2_ segregants that differed at the Chromosome XIV locus and gathered a panel of F_3_ segregants. (D) Linkage mapping in the F_3_ segregants identified the Chromosome XIV locus at high resolution, with a peak at position 467,219. Tick marks denote every 100,000 bases along the chromosome. (E) Recombination breakpoints in the F_3_ segregants delimited the Chromosome XIV locus to a single SNP in *MKT1* at position 467,219. Vertical dashed line highlights the delimited causal polymorphism, while small vertical lines along the x-axis indicate different SNPs in the window that is shown. (F) Engineering the BY allele into *mrp20-105E XIV^3S^* segregants changed growth (left), while substitutions at the nearest upstream and downstream variants did not (right).

To further resolve the Chromosome XIV locus, we crossed an *mrp20-105E XIV^BY^* F_2_ segregant and an *mrp20-105E XIV^3S^* F_2_ segregant (supplementary text 2; Fig. 2C, table S1). 361 F_3_ progeny were genotyped by low-coverage whole genome sequencing and phenotyped for growth. Linkage mapping with these data reidentified the Chromosome XIV locus at a p-value of 2.50 × 10^−43^ (ANOVA; Fig. 2D, fig. S3) and resolved it to a single SNP in the coding region of *MKT1* (Fig. 2E). This SNP, which encodes a glycine in BY and a serine in 3S at amino acid 30, was then validated by nucleotide replacement in *mrp20-105E* segregants (Fig. 2F). Notably, this specific SNP was previously shown to play a role in mitochondrial genome stability (24), suggesting epistasis between *MRP20* and *MKT1* involves mitochondrial dysfunction, impairing growth on non-fermentative carbon sources such as ethanol.

### Epistasis between *MRP20* and *MKT1* differs in cross parents and segregants

We attempted to validate the epistasis between *MRP20* and *MKT1* by introducing all four possible combinations of the causal nucleotides at these two genes into haploid versions of both BY and 3S (Fig. 3A). The *mrp20-105E* mutation affected growth in both parent strains (ANOVA, p = 4.3 × 10^−24^ and p = 4.0 × 10^−4^). However, the magnitude of the effect differed between the two: *mrp20-105E* caused inviability in BY but had a more modest effect in 3S. In addition, *MKT1* influenced response to *mrp20-105E* in the 3S background (ANOVA, p = 0.01) but not in the BY background (ANOVA, p = 0.99).

**Figure 3.**
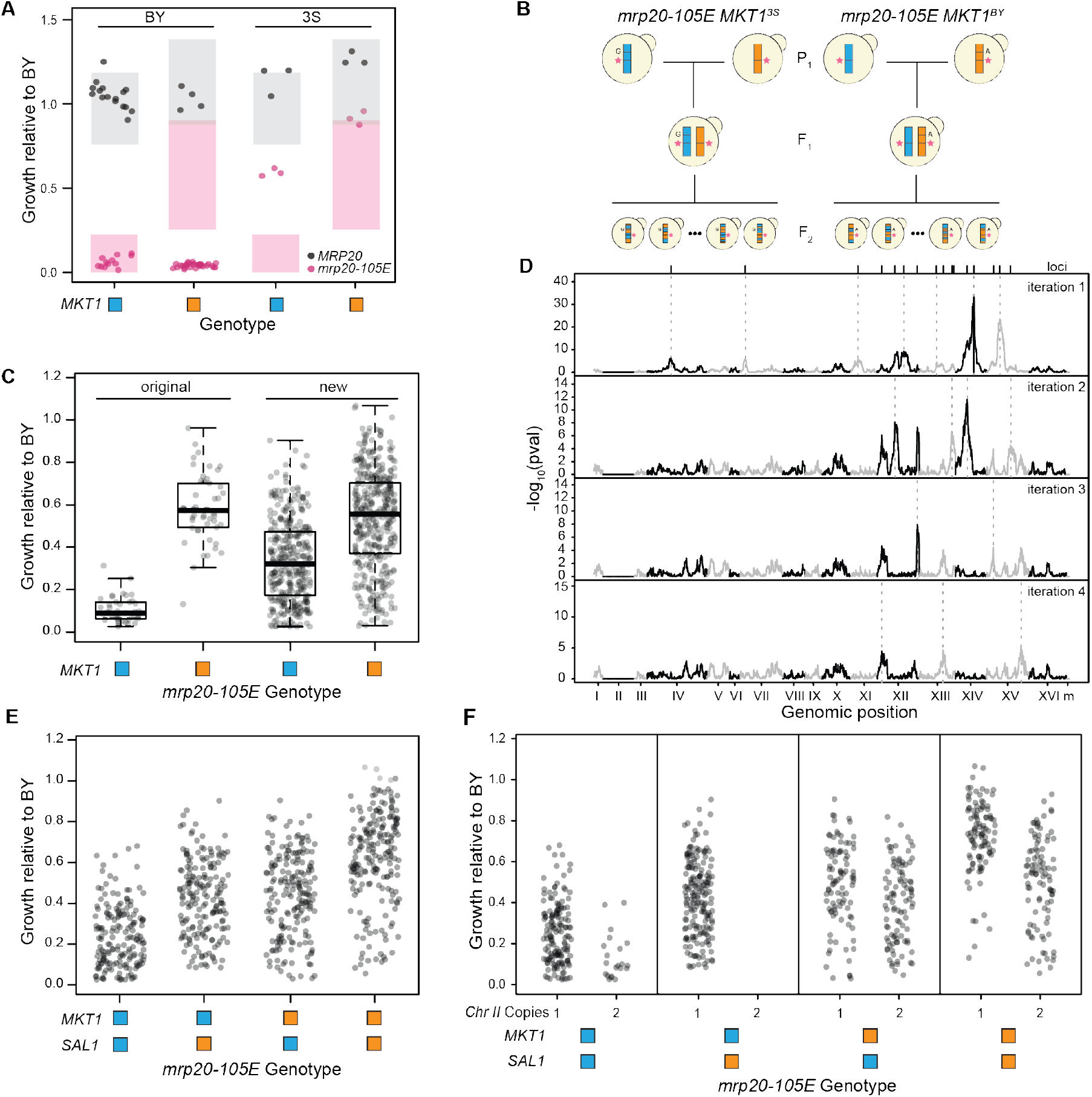
Additional loci govern response to the mutation. (A) We engineered all combinations of *MRP20* and *MKT1* into the BY and 3S cross parents. Expected phenotypes are shown as shaded boxes denoting 95% confidence interval based on the originally obtained segregant phenotypes. (B) We generated BY x 3S crosses in which all segregants carried *mrp20-105E*. Two crosses were performed: one in which all segregants carried *MKT1^BY^* and one in which all segregants carried *MKT1^3S^*. Tetrads were dissected and spores were phenotyped for growth on ethanol. (C) Each of the new crosses showed increased growth that extended from inviable to wild type, differing from the more qualitative bimodal phenotypes seen among the original *mrp20-105E MKT1* segregant populations. (D) Linkage mapping identified a total of 16 loci that influenced growth. After four iterations of a forward regression, no additional loci were identified. (E) Inviable segregants were present among all *mrp20-105E MKT1 SAL1* genotype classes. (F) Aneuploid individuals with duplicated Chromosome II showed reduced growth. Aneuploid individuals were not evenly detected across the different *MKT1*-*SAL1* genotype classes.

The phenotypic consequence of epistasis between *MRP20* and *MKT1* differed between parent and segregant strains. Specifically, the phenotypes of BY *mrp20-105E MKT1^3S^*, 3S *mrp20-105E MKT1^BY^*, and 3S *mrp20-105E MKT1^3S^* all differed from the expectations established by BYx3S *mrp20-105E* segregants. These departures from expectation imply that additional unidentified loci also influence response to *mrp20-105E*.

### Fixation of *mrp20-105E* and *MKT1* genotypes increases phenotypic variance

To enable the identification of other loci underlying response to *mrp20-105E*, we generated two new BYx3S crosses (Fig. 3B, table S1). In both crosses, the BY and 3S parents were engineered to carry *mrp20-105E*. Furthermore, one cross was engineered so that both parents carried *MKT1^BY^* and the other cross was engineered so that both parents carried *MKT1^3S^*. By altering the parent strains in this manner, we increased the chance of detecting additional loci contributing to the variable expressivity of *mrp20-105E*. From these engineered crosses, 749 total segregants were obtained through tetrad dissection, genotyped by low-coverage genome sequencing, and phenotyped for growth on ethanol.

The new crosses exhibited continuous ranges of phenotypes, in contrast to the bimodal phenotypic distribution observed in the original segregants (Fig. 3C). In both the *MKT1^BY^* and *MKT1^3S^* crosses, *mrp20-105E* segregants ranged from inviable to nearly wild type. The distributions of phenotypes in the two crosses differed in a manner consistent with their *MKT1* alleles, with the mean of the *MKT1^BY^* segregants lower than the *MKT1^3S^* segregants (t-test, p = 4.8 × 10^−34^). These data show that regardless of the *MKT1* allele present, additional loci can cause *mrp20-105E* to show phenotypic effects ranging from lethal to benign.

### Many additional loci affect the expressivity of *mrp20-105E*

We used the new crosses to map other loci contributing to response to *mrp20-105E*. Excluding *MKT1*, which explained 18% of the phenotypic variance in the new crosses, linkage mapping identified 16 new loci (Fig. 3D, fig. S4, and table S2). We found no evidence for genetic interactions among the loci (pairs and trios examined with fixed effects linear models, Bonferroni threshold).

Of the new loci, the BY allele was inferior at 10 and superior at six. These loci individually explained between 0.79% and 14% of the phenotypic variance in the new crosses. 13 of these loci resided on a subset of chromosomes but were distantly linked: four on Chromosomes XII, three on XIII, two on XIV, and four on XV. The three remaining loci were detected on Chromosomes IV, VII, and XI.

Recombination breakpoints delimited the loci to small genomic intervals spanning one (12 loci), two (3 loci) or three (1 locus) genes (table S3). These candidate genes functioned in many compartments of the cell and implicated a diversity of cellular pathways and processes in the expressivity of *mrp20-105E* (table S4). Thus, the molecular basis of *mrp20-105E*’s expressivity is complex.

### The Chromosome XIV locus contains multiple causal variants

Among the newly detected loci, the largest effect (14% phenotypic variance explained) was on Chromosome XIV. The position of maximal significance at this site was located two genes away from the end of *MKT1*, with a 99% confidence interval that did not encompass the causal variant in *MKT1* (table S3). Thus, the originally identified large effect Chromosome XIV locus in fact represents multiple distinct closely linked nucleotides that both genetically interact with *MRP20* and occur in different genes (Fig. 3E).

The new locus on Chromosome XIV was delimited to two genes, one of which was *SAL1*, encoding a mitochondrial ADP/ATP transporter that physically interacts with Mrp20. A SNP in *SAL1* that segregates in this cross was previously linked to increased mitochondrial genome instability in BY (24), suggesting it is likely also causal in our study. For this reason, we refer to this additional Chromosome XIV locus as *‘SAL1*’. We found no evidence for epistasis between *MKT1* and *SAL1* (ANOVA, p = 0.77).

Although the *MKT1-SAL1* locus had a large effect, it explained a minority of the phenotypic variance among *mrp20-105E* segregants in a model including all detected loci (32% for *MKT1-SAL1* vs. 36% for all other loci collectively). Thus, by enabling *MKT1* and *SAL1* to segregate independently through genetic engineering and examining a large number of *mrp20-105E* segregants with different *MKT1-SAL1* genotypes, we observed a greater diversity of mutational responses than was originally seen and detected many additional loci.

### Aneuploidy also contributes to the expressivity of *mrp20-105E*

Despite the fact that the identified loci explain most of *mrp20-105E*’s expressivity, some individuals exhibited unexpectedly poor growth (Fig. 3F). This finding led to the identification of a Chromosome II duplication that reduced growth (ANOVA, 1.2 × 10^−48^). The aneuploidy was common among *mrp20-105E* segregants, with a higher prevalence when *MKT1^3S^* was also present (Fisher’s exact test, p = 1.5 × 10^−43^; table S5). The Chromosome II aneuploidy was not seen among wild type segregants. These data suggest that *mrp20-105E* increases the rate of aneuploidization and that genetic variation in *MKT1* influences the degree to which *mrp20-105E* segregants duplicate Chromosome II. The aneuploidy’s contribution to phenotypic variation was relatively minor, explaining 5% of phenotypic variance among *mrp20-105E* segregants in a model also including all identified loci.

### Multiple mechanisms underlie poor growth in the presence of *mrp20-105E*

Evidence suggests mitochondrial genome instability contributes to the variable expressivity of *mrp20-105E*. First, mitochondrial genome instability is known to cause poor growth on non-fermentative carbon sources, such as ethanol (25, 26). Second, the exact variants that segregate in our cross at *MKT1* and *SAL1* were previously linked to mitochondrial genome instability (24). Third, both Mrp20 and Sal1 function in the mitochondria (22, 27). Fourth, two other candidate genes in the newly detected loci encode proteins that function in the mitochondria (table S4).

To determine the role of mitochondrial genome instability in the variable expressivity of *mrp20-105E*, we measured petite formation, a proxy for spontaneous mitochondrial genome loss (Fig. 4) (28). In addition to *MRP20* and *mrp20-105E* BY and 3S parent strains, 16 *MRP20* segregants and 42 *mrp20-105E* segregants were examined. Despite causing reduced growth in both parents, *mrp20-105E* only led to elevated mitochondrial genome instability in BY (t-test p = 0.013 in BY and p = 0.39 in 3S; Fig. 4A). Also, although *mrp20-105E* segregants exhibited increased mitochondrial genome instability relative to *MRP20* segregants (Wilcoxon rank-sum test p = 0.023), especially at lower levels of growth, a subset of inviable segregants did not show elevated petite formation (Fig. 4B and C). These results suggest that mitochondrial genome instability explains part, but not all, of response to *mrp20-105E*.

**Figure 4.**
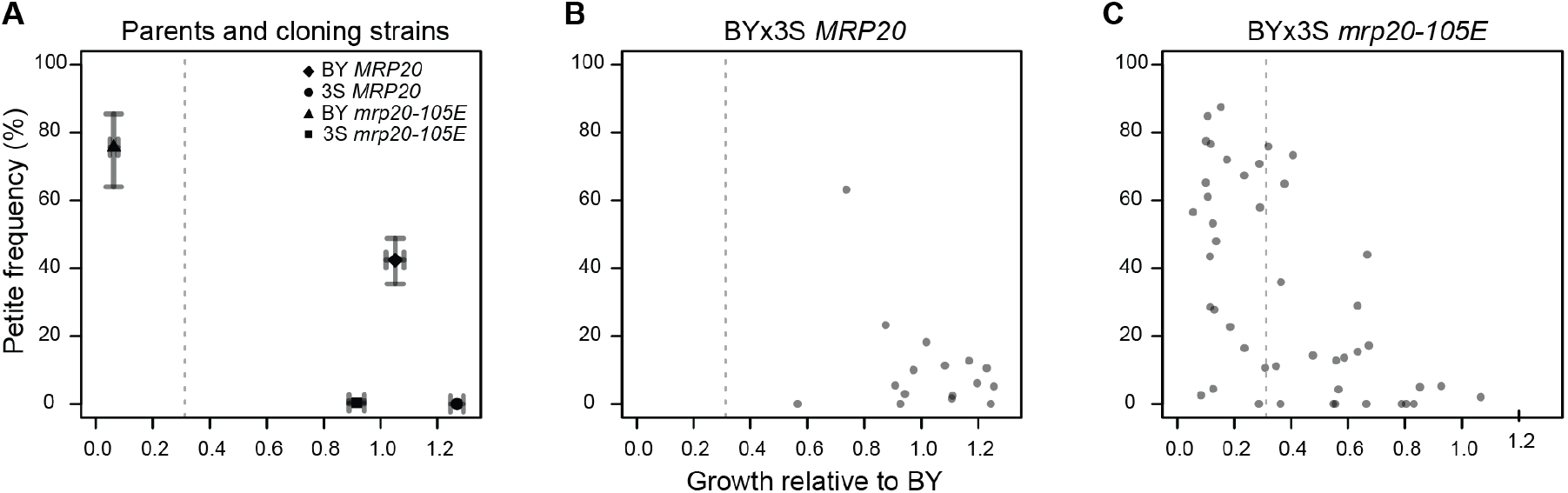
Mitochondrial genome instability partially underlies the expressivity of *mrp20-105E*. We measured petite formation frequency, which estimates the proportion of cells within a clonal population capable of respiratory growth. Higher petite frequency is a proxy for greater mitochondrial genome instability. (A) We examined *MRP20* and *mrp20-105E* versions of the BY and 3S parent strains. For each, average values and 95% bootstrapped confidence intervals are shown. BY showed elevated mitochondrial genome instability in the presence of *mrp20-105E*, while 3S showed no change. (B) We examined 16 BYx3S *MRP20* segregants. These segregants were randomly selected and spanned the range of growth values for *MRP20* segregants. (C) 45 BYx3S *mrp20-105E* segregants. Poorer growing segregants tended to exhibit higher mitochondrial genome instability, though some exhibited wild type levels of mitochondrial genome instability. The gray dashed line indicates the threshold used to call inviability.

### Genetic underpinnings of *mrp20-105E*’s expressivity in segregants and parents

We determined the extent to which our identified loci explained phenotypic variability among mutants. Modeling growth as a function of all identified loci and the aneuploidy accounted for the majority (78%) of the broad-sense heritability among *mrp20-105E* segregants (ANOVA, p = 5.2 × 10^−188^). Further, phenotypic predictions for segregants based on their genotypes were strongly correlated with their observed phenotypes (r = 0.85, p = 4.4 × 10^−209^; Fig. 5A). These results show that the variable expressivity of *mrp20-105E* is driven by many loci that collectively produce a spectrum of mutational responses.

**Figure 5.**
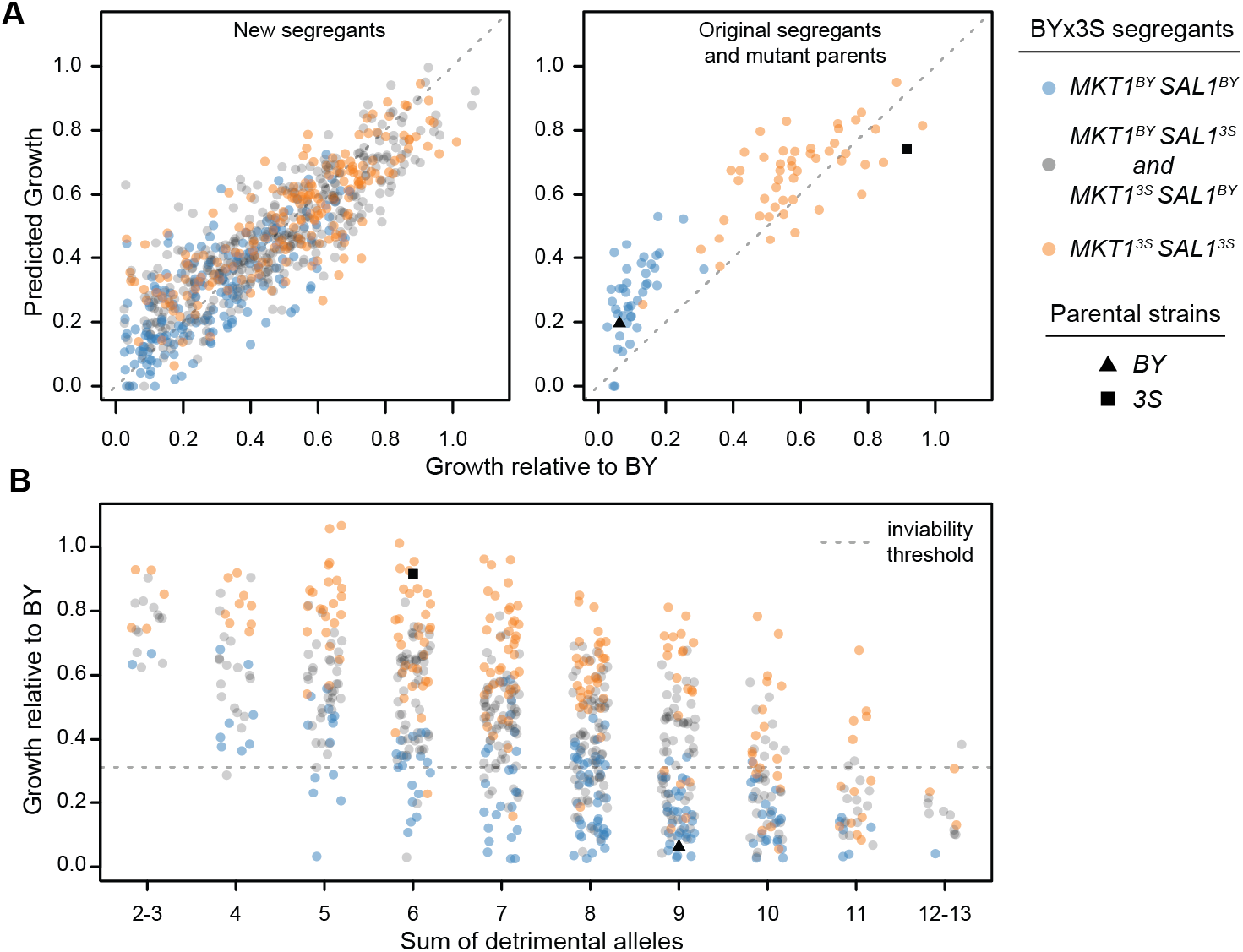
Detected loci quantitatively and qualitatively explain mutant phenotypes. (A) We fit a linear model accounting for the effects of all detected loci and the aneuploidy on the growth of *mrp20-105E* segregants. This model not only explained the growth of the new BY x 3S *mrp20-105E* crosses generated in this paper, but also accurately predicted the phenotypes of the mutant parents and previously generated segregants. (B) We examined growth relative to the sum of detrimental alleles carried by a segregant. This relationship shows how collections of loci produce a quantitative spectrum of phenotypes, including instances of qualitative phenotypic responses. This relationship explains the full range of responses, from inviable to wild type growth, across *MKT1-SAL1* genotypes. The gray dashed line indicates the threshold used to call inviability.

Confirming this point, the model was also effective for other genotypes that were not present in the new crosses, but had been generated throughout the course of this work. For instance, the model accurately predicted the phenotypes of the original *mrp20-105E* segregant population (r = 0.90, p = 1.6 × 10^−39^), as well as the phenotypes of cross parents engineered to carry *mrp20-105E* (Fig. 5A). Moreover, the model explained both qualitative and quantitative variation within and between the two Chromosome XIV classes that were originally seen among *mrp20-105E* segregants.

Finally, we examined how diverse combinations of loci collectively produced similar phenotypic responses to *mrp20-105E*. We examined the relationship between growth and the total number of detrimental alleles carried by *mrp20-105E* segregants, keeping track of each individual’s genotype at *MKT1* and *SAL1*, the largest effect loci (Fig. 5B). The number of detrimental alleles carried by a segregant showed a strong negative relationship with growth, which was not observed in wild type segregants (fig. S5). Further, regardless of genotype at *MKT1* and *SAL1*, the effect of *mrp20-105E* ranged from lethal to benign in a manner dependent on the number of detrimental alleles present at other loci. These findings demonstrate that many segregating loci beyond the large effect *MKT1-SAL1* locus influence the expressivity of *mrp20-105E* and enable different genotypes in the cross to exhibit a broad range of responses to the mutation.

## Discussion

We have provided a detailed genetic characterization of the expressivity of a spontaneous mutation. Response to this mutation in a budding yeast cross is influenced by at least 18 genetic factors in total, with the largest effect due to two closely linked variants. However, at least 15 additional loci segregate and jointly exert larger effects than the largest two. Different combinations of alleles across these loci produce a continuous spectrum of mutational responses. Due to tight linkage between *MKT1* and *SAL1* in the original cross parents, the full extent of this continuum was not originally observed, leading to an initial understanding of the expressivity of the *mrp20-105E* mutation that was simplistic.

These findings also show how quantitative variation in mutational response can produce seemingly discrete outcomes. In part, whether responses appear qualitative depends on the configuration of mutationally responsive alleles in examined mutants. Approaches such as crossing of genetically engineered strains can be used to disrupt these configurations that mask the full extent of variation. However, another part of this expressivity is the tolerance of a system to quantitative variation in key processes, for example mitochondrial genome stability in the case of *mrp20-105E*. Our data suggest that these processes can only tolerate quantitative variation to a point, but also indicate that lethality to the same mutation may arise in different genetic backgrounds due to impairment of distinct cellular processes.

Our results inform efforts to understand expressivity in other systems, including humans. For example, there is interest in determining why people who carry highly penetrant alleles known to cause disease do not develop pathological conditions (3, 29, 30). Such resilience, as observed here, may involve numerous loci. This speaks to the complicated and unexpected epistasis that can arise between mutations and segregating loci in genetically diverse populations (7–20). It also illustrates the importance of characterizing epistasis (31–41), including background effects, as these forms of genetic interactions are immediately relevant to evolution and disease, and may not emerge from studies that do not directly interrogate natural variation in genetically diverse populations.

## Materials and Methods

### Generation of segregants

The haploid BYx3S segregants in which *mrp20-105E* was identified were the *hos3*Δ F_2_ segregants generated and described in Mullis et al (17). In brief, a BY *MATa can1*Δ::*STE2pr-SpHIS5 his3*Δ strain was mated to a 3S *MATα ho*::*HphMX his3*Δ::*NatMX* strain to generate a wild type BY/3S diploid. PCR-mediated, targeted gene disruption was then used to produce a BY/3S *HOS3/hos3*Δ::*KanMX* strain. Both the wild type and hemizygous deletion strains were sporulated, and random BYx3S *MATa* spores were obtained from each using the magic marker system with plating on His- plates containing canavanine (42). Following discovery of the *mrp20-105E* mutation, we performed tetrad dissected of this diploid to obtain *mrp20-105E* segregants in both *HOS3* and *hos3*Δ genetic backgrounds.

To produce haploid *mrp20-105E* F_3_ segregants, we deleted *URA3* from a BYx3S F_2_ *MATa can1*Δ::*STE2pr-SpHIS5 his3*Δ *hos3*Δ::*KanMX mrp20-105E XIV^3S^* segregant. We then mating type switched the strain by transforming it with a *URA3* plasmid containing an inducible *HO* endonuclease, inducing *HO*, and plating single cells. The mating type-switched BYx3S F_2_ *MATα can1*Δ::*STE2pr-SpHIS5 his3*Δ *ura3*Δ *hos3*Δ::*KanMX mrp20-105E XIV^3S^* segregant was then mated to a BYx3S F_2_ *MATa can1*Δ::*STE2pr-SpHIS5 his3*Δ *hos3*Δ::*KanMX mrp20-105E XIV^BY^* segregant. The resulting diploid was sporulated and random segregants were obtained by plating on His- media.

To obtain additional haploid *mrp20-105E MKT1^BY^* and *MKT1^3S^* F_2_ segregants, we engineered *mrp20-105E*, as well as the 3S and BY causal variants at *MKT1* position 467,219 into BY and 3S, respectively. BY *mrp20-105E* was independently mated to 3S *mrp20-105E MKT1^BY^* twice. Two resultant diploids were sporulated and tetrads were dissected to obtain BYx3S *mrp20-105E MKT1^BY^* haploid segregants. The same process was followed with BY mrp20-105E *MKT1^3S^* and 3S *mrp20-105E* strains to obtain BYx3S *mrp20-105E MKT1^3S^* haploid segregants.

### Genotyping

F_2_ segregants shown in Fig 1 and Fig 2 A-B were previously genotyped in Mullis et al. using the same techniques described below (17). In this paper, F_3_ segregants and all remaining F_2_ segregants shown in Figs 1 and 3–5 were genotyped by low coverage whole genome sequencing. Freezer stocks of strains were inoculated into liquid overnight cultures and grown to stationary phase at 30°C. DNA was extracted using Qiagen 96-well DNeasy kits (Qiagen P/N 69581). Sequencing libraries were prepared using the Illumina Nextera Kit and custom barcoded adapter sequences. Segregants from each respective cross (361 F_3_s and 872 F_2_s) were pooled in equimolar fractions into three separate multiplexes, run on a gel, size selected, and purified with the Qiagen Gel Extraction Kit. F_2_ and F_3_ segregants were sequenced by Novogene on Illumina HiSeq 4000 lanes using 150 bp x 150 bp paired-end reads.

Sequencing reads were mapped against the S288C genome (version S288C_reference_sequence_R64-2-1_20150113.fsa from the Saccharomyces Genome Database https://www.yeastgenome.org) using BWA version 0.7.7-r44 (43). Samtools v1.9 was then used to create a pileup file for each segregant (44). For both BWA and Samtools, default settings were employed. Base calls and coverages were gathered for 44,429 SNPs that segregate in the cross (14). Low coverage individuals (<0.7x average per site coverage) were removed from analyses. Diploid and contaminated individuals were identified by abnormal patterns of heterozygosity or sequencing coverage, and were also excluded. For each segregant, a raw genotype vector was determined by the percent of calls at each site for the 3S allele. We then used a Hidden Markov Model (HMM) implemented in the ‘HMM’ package v 1.0 in R to correct each raw genotype vector using the following probability matrices (45): transitionProbabilitiy = matrix(c(.9999,.0001,.0001,.9999),2) and emissionProbability = matrix(c(.0.25,0.75,0.75,0.25),2).

Aneuploidies were identified based on elevated sequencing coverage at particular chromosomes within each individual sample. This identified a chromosome II duplication event in a subset of BYx3S *mrp20-105E MKT1^BY^* and BYx3S *mrp20-105E MKT1^3S^* segregants. The BY *mrp20-105E MKT1^3S^* x 3S *mrp20-105E* cross had the highest prevalence (50%), and thus individuals from this cross were further examined. We employed the normalmixEM() function from the mixtools library in R (46) to determine that coverage on Chr II was bimodal and centered on 0.98 and 1.8 (log likelihood of 237). Posterior probabilities were used to call aneuploid individuals which that had an average per site coverage of 1.5x or greater. This threshold was also applied to other crosses to identify aneuploid individuals.

### Phenotyping

Segregants were inoculated into rich media containing glucose (‘YPD’), which was comprised of 1% yeast extract (BD P/N 212750), 2% peptone (BD P/N: 211677), and 2% dextrose (BD P/N 15530). Cultures were grown to stationary phase (two days at 30°C). Strains were then pinned onto YP + 2% agar (BD P/N 214050) rich media containing ethanol (‘YPE’). The YPE recipe was 1% yeast extract (BD P/N 212750), 2% peptone (BD P/N: 211677), and 2% ethanol (Koptec P/N A06141602W). Plates were then grown at 30°C for two days. Growth assays were conducted in a minimum of three replicates across three plates. On each plate, a BY control was included. Plates were imaged with the BioRAD Gel Doc XR+ Molecular Imager at a standard size of 11.4 × 8.52 cm^2^ (width x length) and imaged with epi-illumination using an exposure time of 0.5 seconds. Images were saved as 600 dpi tiffs. ImageJ (http://rsbweb.nih.gov/ij/) was used to quantify pixel intensity of each colony through the Plate Analysis JRU v1 plugin (https://research.stowers.org/imagejplugins/zipped_plugins.html), as described in Matsui et al. (47). Growth values were normalized against the same plate BY control, then averaged across replicates to produce a single growth value for each segregant.

### Linkage mapping

Initial linkage mapping was conducted with F_2_ segregants. Initial discovery of the spontaneous *mrp20-105E* mutation resulted from linkage mapping with 385 F_2_ segregants (164 wild type and 221 *hos3*Δ) from Mullis et al. (17). We employed the linear model *growth* ~ *hos3*Δ *+ locus + hos3*Δx*locus + error*, from which the *hos3*Δx*locus* interaction term was used to identify loci that differentially explained growth in *hos3*Δ segregants. Examination of the *hos3*Δx*locus* interaction term led to discovery of the spontaneous *mrp20-105E* mutation on *the MRP20^BY^* allele present in *hos3*Δ segregants. Following discovery of *mrp20-105E*, we used the fixed effects linear model *growth ~ MRP20 + locus + MRP20 x locus + error* using only *hos3*Δ individuals from Mullis et al. (17). From this scan, we examined the *MRP20* x *locus* interaction term. 361 *mrp20-105E* F_3_ segregants were used to better resolve the Chromosome XIV locus. We employed the model *growth* ~ *locus* + *error* and examined the *locus* term. We examined the minimum observed test on chromosome XIV to delimit that locus.

To find loci affecting growth in the *mrp20-105E* background, we generated new populations of *mrp20-105E MKT1^BY^* (353) and *mrp20-105E MKT1^3S^* (396) haploid segregants. The combined 749 *mrp20-105E* segregants were used for linkage mapping that followed a forward regression approach. We first obtained residuals from the linear model *growth ~ MKT1 + error*, and then implemented a genome-wide scan using the model *residuals ~ locus + error*. We examined the locus term and significance was determined by using 1,000 permutations with the threshold set at the 95^th^ quantile of observed −log_10_(p-values) (48). A maximum of one locus per chromosome per scan was identified as significant. Following the identification of additional loci, we accounted for the newly detected loci in a new model, *residuals ~ locus 1 + locus 2 + … locus n + error* and obtained the residuals. These new residuals were used in another genome-wide scan using the model *residuals ~ locus + error*. Permuted thresholds were calculated for each scan. This process was repeated for a total of 5 iterations at which point no loci were detected above our significance threshold. Chromosome II was excluded from linkage mapping due to the presence of a chromosomal duplication in a subset of individuals. The Chromosome II duplication was tested for significance using the model *growth ~ MKT1 + ChromosomeII + error,* from which the Chromosome II term was examined.

All linkage mapping was performed in R. Linear models were implemented using the lm() function. To call peaks for each scan we required that the local minimum position within each peak be a minimum of 150,000 kb away from any other peak. We also required peaks to be more than 20kB from the edge of a chromosome. We report 99% confidence intervals as 2-lod intervals surrounding the peak position at each locus.

### Classification of inviable segregants

Initial discovery of the *MRP20 x MKT1* genetic interaction suggested that expressivity of *mrp20-105E* was largely determined by variation at *MKT1*. Furthermore, *mrp20-105E MKT1^BY^* segregants exhibited very poor growth, while *mrp20-105E MKT1^3S^* segregants showed more tolerant, variable growth. We termed this initial *mrp20-105E MKT1^BY^* segregant population as ‘inviable’. Figures 4 and 5 include a gray dashed line to denote the highest growth value observed among the original inviable segregants.

### Delimiting loci with recombination breakpoints

For each locus examined, we split the appropriate segregants into two groups: individuals carrying the BY allele and individuals carrying the 3S allele. Segregants’ haplotypes across the adjacent genomic window were then examined. The causal region was determined by identifying the SNPs fixed for BY among all BY individuals and fixed for 3S among all 3S individuals. Raw Illumina sequencing reads were examined to confirm the delimit of *IV* to *MRP20* among original F_2_ segregants, the delimit of *XIV* to the *MKT1* coding SNP at 467,219 among F_3_ segregants, and the delimit of the secondary *XIV* locus to *SAL1* and *PMS1* among the new F_2_ segregants.

### Reciprocal hemizygosity experiments

Four *hos3*Δ F_2_ *MATa* segregants were used in all reciprocal hemizygosity (RH) experiments (49): two were *hos3Δ IV^BY^ XIV^BY^* and two were *hos3Δ IV^3S^ XIV^BY^*. The four segregants were first mating type switched to enable mating of these segregants to produce homozygous *IV^BY^/IV^BY^*, homozygous *IV^3S^/IV^3S^*, or heterozygous *IV^BY^/IV^3S^* diploids. Each pairwise mating was performed and confirmed by plating on mating type tester plates. These diploid strains were then phenotyped on agar plates containing ethanol, which verified that *IV^BY^* has an effect in diploids and acts in a recessive manner. Using the haploid *MATa* and *MATα* versions of these four segregants, we individually engineered premature stop codons into *DIT1*, MRP20 and *PDR15* using CRISPR-mediated targeted gene disruption and lithium acetate transformations (50). Plasmid-based CRISPR-Cas9 was employed to target the beginning of each coding region and 20bp repair templates which contained a premature stop codon followed by 1bp deletions were incorporated. Each sgRNA and repair template was designed so that only the first 15 (of 537), 26 (of 264), and 33 (of 1,530) amino acids would be translated for *DIT1, MRP20* and *PDR15*, respectively. Engineered strains were confirmed by PCR and Sanger sequencing. After confirmation, wild type and knockout strains for each gene were then mated in particular combinations to produce reciprocal hemizygotes that were otherwise isogenic. A minimum of two distinct hemizygotes were generated for each allele of each gene.

### Construction of nucleotide replacement strains

Single nucleotide replacement strains were generated for *MRP20* and *MKT1* using a CRISPR/Cas9-mediated 10pproach. For a given replacement, the appropriate strain was first transformed with a modified version of pML104 that constitutively expresses Cas9 using LiAc transformation (50, 51). We then inserted the KanMX gene using co-transformation of a double-stranded DNA containing KanMX with 30bp upstream and 30bp downstream homology tails and gRNAs targeting the region containing the site of interest (52). DNA oligos and PCR were used to construct custom sgDNA templates which included crRNA and tracrRNA in a single molecule. Next, we employed T7 RNA Polymerase to express sgDNA templates *in vitro*. Dnase treatment and phenol extraction were used to obtain purified sgRNAs. Transformants were selected on media containing G418, and KanMX integration was confirmed by PCR. Next, KanMX was replaced with the nucleotide of interest. To do this, integrants were co-transformed with four gRNAs targeting KanMX, a 60 bp single-stranded DNA repair oligo, and a marker plasmid expressing either HygMX or NatMX using electroporation (53). Marker plasmids were constructed by Gibson assembly with HygMX or NatMX and pRS316 (54, 55). Repair constructs were 60bp ssDNA oligos ordered from Integrated DNA technologies that included upstream homology, the desired nucleotide at the site of interest, and downstream homology. Transformants were selected on media containing either hygromycin or nourseothricin, depending on what marker plasmid was used. Replacement strains were then confirmed by sanger sequencing.

Following this strategy, *the mrp20-105E* nucleotide was engineered into two *hos3Δ IV^3S^ XIV^BY^* segregants, and two *hos3Δ IV^BY^ XIV^BY^* segregants were restored to *MRP20*. Similarly, at *MKT1* the causal, nearest upstream and downstream SNPs were engineered into two *hos3Δ IV^BY^ XIV^3S^* segregants. Similarly, we generate BY *mrp20-105E*, BY *MKT1^3S^*, 3S *mrp20-105E*, and 3S *MKT1^BY^* strains in this manner. Each single nucleotide parental replacement strain was then backcrossed to its own progenitor. Each subsequent diploid was sporulated and tetrad dissected, and we confirmed haploid genotypes by sequencing. The same approach was used to generate 3S mrp20-105E *MKT1^BY^* haploids by crossing 3S *mrp20-105E* and 3S *MKT1^BY^* strains. However, this strategy could not be followed to generate BY *mrp20-105E MKT1^3S^ h*aploids, because, crossing BY *mrp20-105E* and BY *MKT1^3s^* strains failed to produce any tetrads with 4 viable spores. Instead, we took BY *MKT1^3S^* strains and converted *MRP20* to *mrp20-105E*.

### Mitochondrial genome instability experiments

We performed *petite* frequency assays as described in Dimitrov et al. (24) In brief, freezer stocks were streaked onto solid YPD media and grown for two days at 30°C. Single colonies were then resuspended in PBS, plated across dilutions onto YPDG plates (1% yeast extract, 2% peptone, 0.1% glucose, and 3% glycerol) and grown for five days at 30°C. Plates were then imaged with the BioRAD Gel Doc XR+ Molecular Imager at a standard size of 12.4 × 8.9 cm^2^ (width x length) and imaged with epi-illumination using an exposure time of 0.5 seconds. Images were saved as 600 dpi tiffs. ImageJ (http://rsbweb.nih.gov/ij/) was used to examine growth and quantitate colony size as described in Dimitrov et al. (24). Colonies were then classified as *petite* and *grande* using a threshold defined as the maximum colony diameter of observed *petites* among BY and 3S wild type strains. *Petite* frequency is the ratio of small colonies to total colonies.

### Modeling growth and examining the model in additional segregant populations

We modeled growth for *mrp20-105E* segregants from the Byx3S crosses fixed for *mrp20-105E* and engineered at *MKT1*. We incorporated MKT1, the 16 detected loci and the Chromosome II duplication in the linear model *growth ~ MKT1 + locus1 + locus2 + … locus16 + Chromsome II + error*. This model was used to generate predicted growth values. We then compared our observed growth values to these predictions. Next, we sought to determine whether loci influencing the expressivity of *mrp20-105E* also affected growth in other strains. To accomplish this, we input the genotype information for each strain into our model to obtain predictions for its growth. We then compared the predicted values to the observed growth values and obtained Pearson correlations when possible.

### Relationship between detrimental alleles, growth, and inviability

At each detected locus influencing response to *mrp20-105E*, we determined the allele associated with worse growth (‘detrimental allele’). Next, we counted the number of detrimental alleles carried by each *mrp20-105E* segregant and examined how phenotypic response to *mrp20-105E* related to it. The *MKT1* and *SAL1* loci were not included when counting detrimental alleles, so that this relationship could be examined across different *MKT1-SAL1* genotype classes.

## Acknowledgments

We thank A. Dudley and G. Cromie for input at an early stage of this project, A. Mahableshwarkar and J. Sloan for assistance with experiments, and A. Coradini, I. Goldstein, and C. Hull for feedback during the implementation and writing phases of this work. We also thank the anonymous reviewers of this manuscript for their valuable feedback and Steven Finkel and his lab for allowing use of laboratory equipment.

## Data Availability Statement

All raw and processed data used in this work is publicly available. Processed data tables and all code used for analyses are included in supplementary Data S1-9 and will be uploaded to the Ehrenreich Lab GitHub page. Raw sequencing data will be available through Bioproject accession PRJNA739014 in the NCBI Short Read Archive.

## Supplementary Information Text

### Resolution of the Chromosome IV locus

We detected a locus on Chromosome IV with a peak marker position from 1,277,231 to 1,277,378, and a 99% confidence interval from position 1,277,231to position 1,278,618, encompassing the promoter and most of the coding region of *MRP20*.

### Resolution of the Chromosome XIV locus

The chromosome XIV locus that interacts with *MRP20* in *hos3*Δ segregants had a peak from 463,554 to 465,005, and a 99% confidence interval that extended from 457,243 to 478,701. 10 protein-coding genes were completely or partially encompassed in this confidence interval limiting possible insight into the causal variation underlying this effect.

**Fig. S1.**
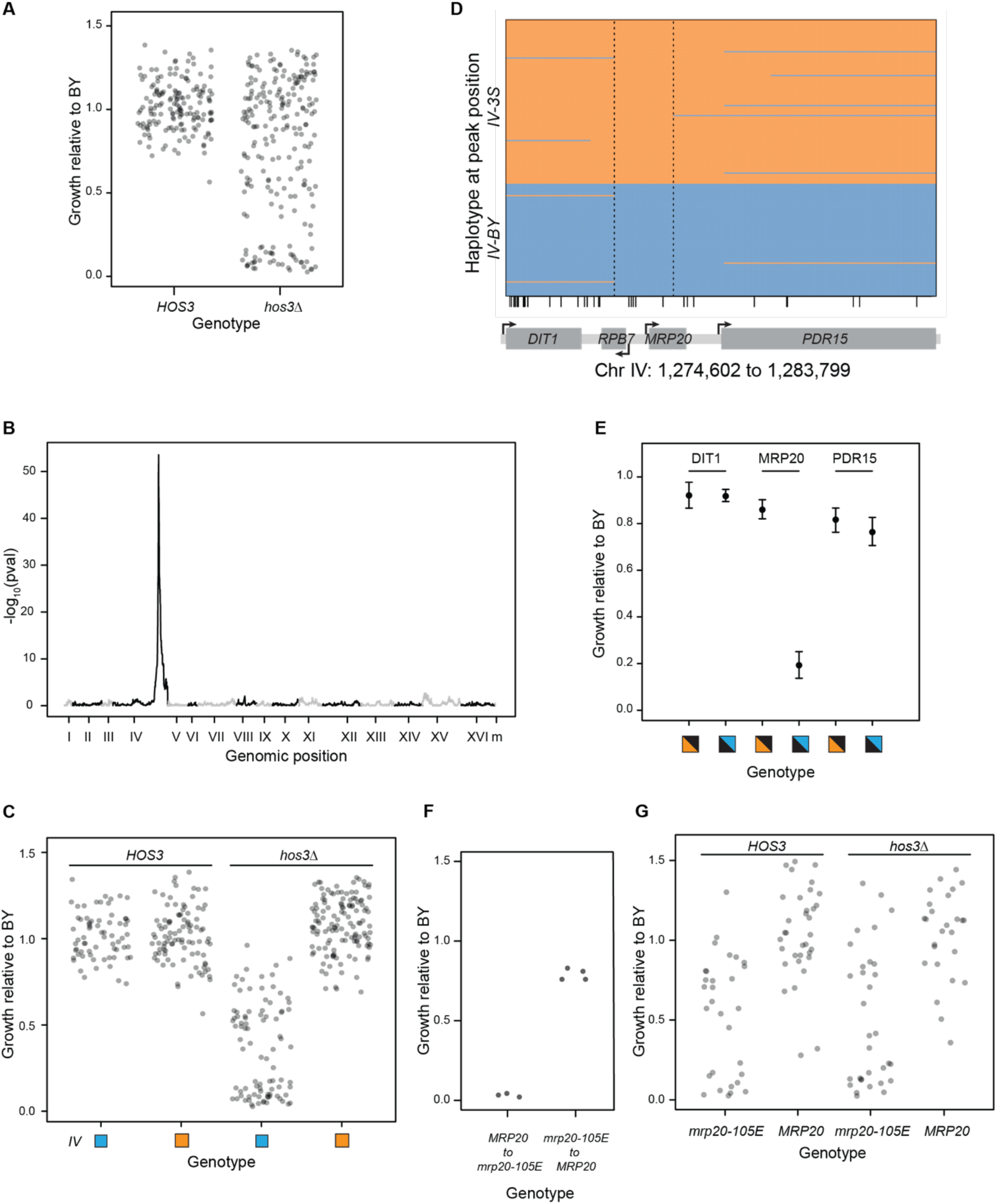
Identification of *mrp20-105E*. (A) Wild type and hemizygous BY/3S diploids were generated and sporulated to produce *HOS3* and *hos3*Δ F2 BYx3S segregants. BYx3S *hos3*Δ segregants exhibited a large increase in phenotypic variability relative to wild type segregants. (B) Linkage mapping using the *HOS3* and *hos3*Δ segregants identified a single locus on Chromosome IV. The peak marker was from 1,277,231 to 1,277,959 and the confidence interval extended from position 1,272,164 to position 1,278,407, encompassing (from left to right) part of *URH1* and all of *DIT2*, *DIT1*, *RPB7*, and *MRP20*. (C) The BY allele of the Chromosome IV locus had a large effect in *hos3*Δ segregants, but no effect in *HOS3* segregants. (D) Recombination breakpoints in *hos3*Δ segregants delimited the Chromosome IV locus to five SNPs (small vertical black lines along the x-axis) in the *RPB7-MRP20* region of the chromosome. Dashed vertical lines show the window delimited by the recombination breakpoints. One of these variants was a spontaneous mutation in *MRP20*. Blue and orange respectively refer to the BY and 3S alleles of the locus. (E) Reciprocal hemizygosity analysis in a *hos3*Δ BY/3S diploid was conducted at closely linked non-essential genes and found that *MRP20* is the causal gene underlying the Chromosome IV locus. In these experiments the *IV^BY^* allele includes the *mrp20-105E* mutation and results in a substantial decrease in growth. Black triangles denote the absence of one allele and colored triangles indicate the alleles that are present. (F) The causality of *mrp20-105E* was validated by engineering in segregants with *MRP20* (left) and *mrp20-105E* (right). (G) Tetrad dissection of the original BY/3S *HOS3/hos3Δ MRP20/mrp20-105E* diploid showed that increased variation was due to *mrp20-105E*, not *hos3*Δ. Throughout the paper, blue and orange are used to denote BY and 3S genetic material, respectively. All growth data presented in the paper are measurements of colonies on agar plates containing rich medium with ethanol as the carbon source.

**Fig. S2.**
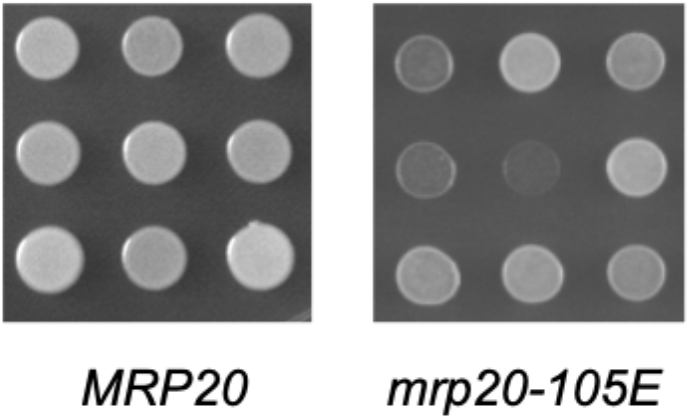
Representative *MRP20* and *mrp20-105E* segregants on ethanol. Each colony is a genetically distinct BYx3S segregant grown on ethanol. A wide range of growth phenotypes was observed among *mrp20-105E* segregants, some of which were inviable in this condition.

**Fig. S3.**
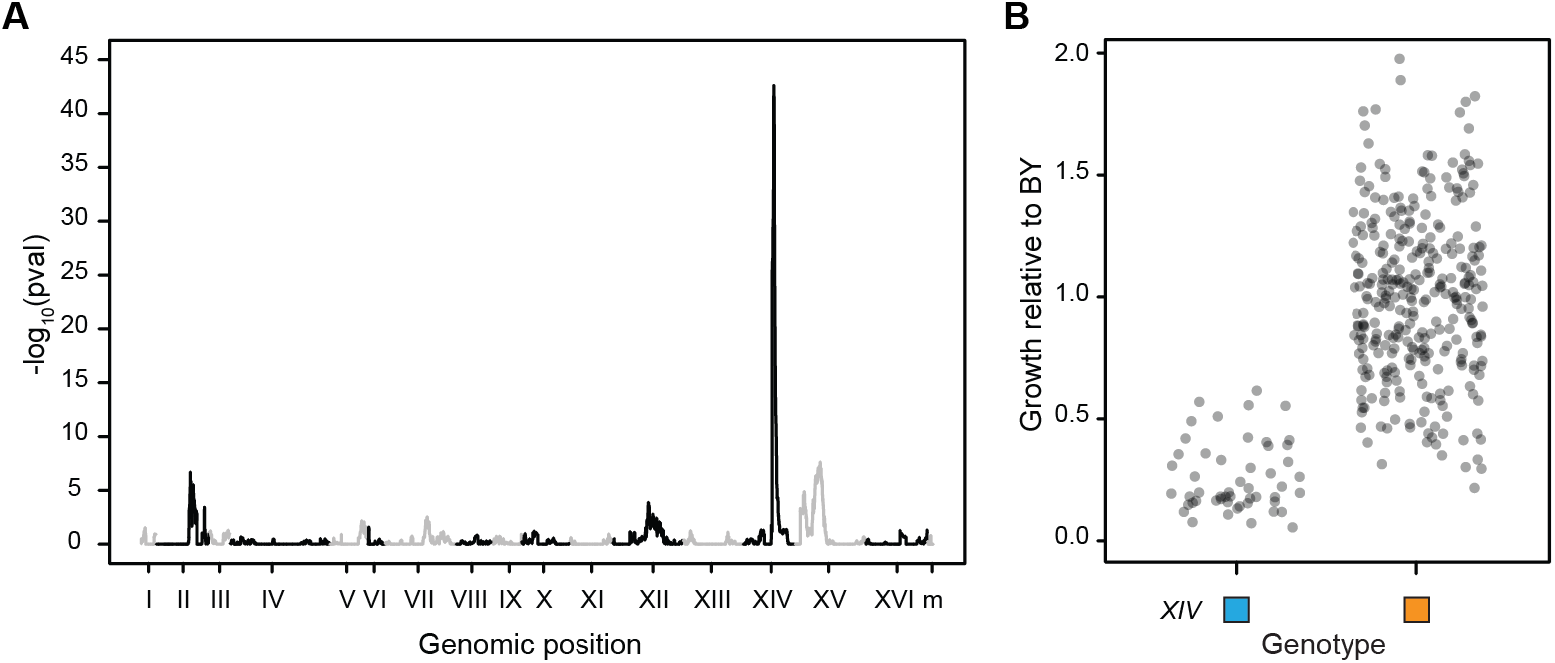
Linkage mapping in the F_3_ panel more finely resolves the Chromosome XIV locus. The model *growth ~ locus + error* was used. The genome-wide significance plot of the *locus* term is shown in (**A**) and the relationship between genotype at the Chromosome XIV locus are shown in (**B**). The peak and 99% confidence interval solely included the position 467,219.

**Fig. S4.**
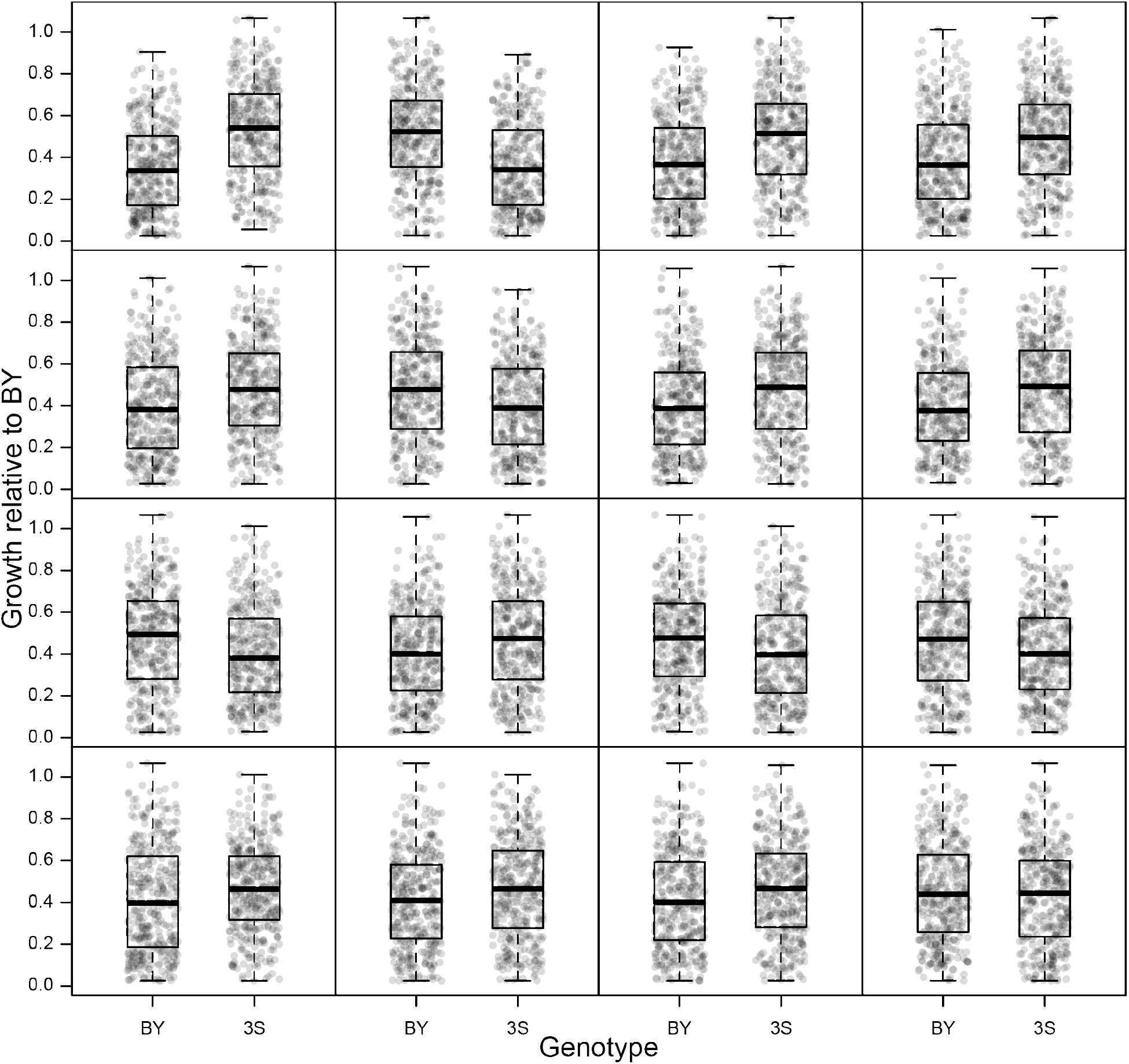
Growth effects of loci detected in BY x 3S *mrp20-105E* crosses. The relationship between genotype is shown at each of the 16 loci detected among BYx3S *mrp20-015E* segregants shown in Fig. 3 B-D. Effects are shown from greatest to least effect size, left to right, top to bottom.

**Fig. S5.**
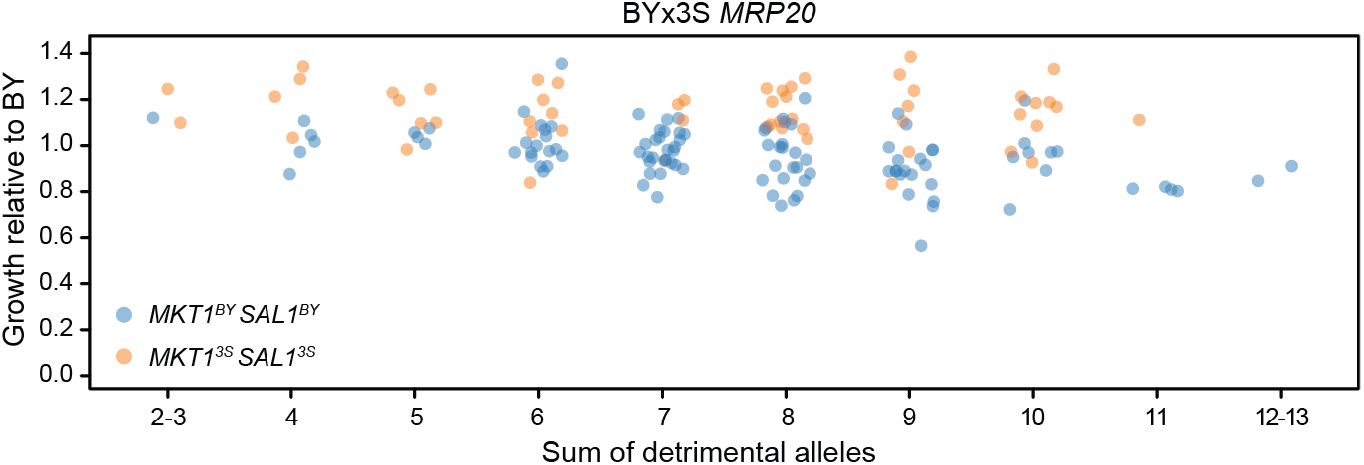
Loci affecting expressivity of *mrp20-015E* show minimal effects in *MRP20* segregants. Growth relative to the sum of detrimental alleles is shown for *MRP20* segregants. While predictions for *MRP20* segregants correlated with observed growth (r = 0.70, p = 9.6 × 10^−25^), the cumulative effects of loci differed between *mrp20-105E* and *MRP20* segregants (ANOVA, observedGrowth ~ predictedGrowth\**MRP20*; interaction term p = 2.8 × 10^−23^). This is likely, in part, due to the fact that wildtype segregants exhibited a narrower range of phenotypes which did not include inviable segregants.

**Table S1.** Crosses and segregant populations examined in this study. All BY x 3S crosses and segregant populations examined in this study are listed. Note, at times different methods of obtaining segregants, either random spores or tetrad dissection were employed.

| Diploid | Cross | Segregants | Method | Publication | Total |
| --- | --- | --- | --- | --- | --- |
| A | BY x 3S | Wildtype F<sub>2</sub> | random spores | Mullis, et al, 2019 | 164 |
| B | BY *mrp20-105E hos3Δ* x<br>3S *MRP20 HOS3* | *MRP20 hos3Δ*<br>F<sub>2</sub> | random spores | Mullis, et al, 2019 | 131 |
|  |  | *mrp20-105E*<br>*hos3Δ* F<sub>2</sub> |  | Mullis, et al, 2019 | 90 |
|  |  | *MRP20 hos3Δ*<br>F<sub>2</sub> | tetrad dissection | this paper | 27 |
|  |  | *mrp20-105E*<br>*hos3Δ* F<sub>2</sub> |  | this paper | 32 |
|  |  | *MRP20 HOS3*<br>F<sub>2</sub> |  | this paper | 34 |
|  |  | *mrp20-105E*<br>*HOS3* F<sub>2</sub> |  | this paper | 30 |
| C | BYx3S *mrp20-105E XIV<sup>BY</sup>* F<sub>2</sub> x<br>BYx3S *mrp20-105E XIV<sup>3S</sup>* F<sub>2</sub> | *mrp20-105E* F<sub>3</sub> | random spores | this paper | 361 |
| D | BY *mrp20-105E MKT1<sup>BY</sup>* x<br>3S *mrp20-105E MKT1<sup>BY</sup>* | *mrp20-105E*<br>*MKT1<sup>BY</sup>* F<sub>2</sub> | tetrad dissection | this paper | 353 |
| E | BY *mrp20-105E MKT1<sup>3S</sup>* x<br>3S *mrp20-105E MKT1<sup>3S</sup>* | *mrp20-105E*<br>*MKT1<sup>3S</sup>* F<sub>2</sub> | tetrad dissection | this paper | 396 |

**Table S2.** Loci beyond *MKT1* that additively influence growth in *mrp20-105E* segregants. These loci were detected by mapping growth ~ locus in in *mrp20-105E MKT^BY^* F_2_ and in *mrp20-105E MKT^3S^* F_2_ individuals shown in Fig 3–5 and Fig S4.

| Chromosome | Peak Position(s) | 99% Confidence Interval | P-value |
| --- | --- | --- | --- |
| 4 | 615,114 | 565,313 to 639,037 | 6.6 x 10<sup>-7</sup> |
| 7 | 140,806 to 142,398 | 86,297 to 164,025 | 1.4 x 10<sup>-6</sup> |
| 11 | 171,261 to 171,457 | 150,554 to 264,387 | 2.1 x 10<sup>-6</sup> |
| 12 | 448,368 | 444,569 to 513,765 | 7.5 x 10<sup>-9</sup> |
| 12 | 679,600 | 660,371 to 701,793 | 4.8 x 10<sup>-10</sup> |
| 12 | 116,314 116,980 | 85,385 to 141,900 | 3.2 x 10<sup>-5</sup> |
| 12 | 1,022,895 to 1,022,933 | 1,013,592 to 1,059,611 | 1.1 x 10<sup>-8</sup> |
| 13 | 612,355 | 594,280 to 652,003 | 2.8 x 10<sup>-5</sup> |
| 13 | 833,534 | 812,150 to 892,748 | 4.6x10<sup>-7</sup> |
| 13 | 449,844 to 451,590 | 436,342 to 465,619 | 3.2 x 10<sup>-5</sup> |
| 14 | 299,515 | 295,050 to 312,454 | 2.0 x 10<sup>-12</sup> |
| 14 | 473,648 | 468,488 to 478,701 | 4.8 x 10<sup>-34</sup> |
| 15 | 185,793 | 167,270 to 189,700 | 3.4 x 10<sup>-5</sup> |
| 15 | 343,484 to 343,921 | 340,625 to 363,553 | 2.4 x 10<sup>-24</sup> |
| 15 | 627,315 to 628,209 | 553,072 to 717,181 | 2.7 x 10<sup>-5</sup> |
| 15 | 885,437 to 885,914 | 869,837 to 916,094 | 5.0 x 10<sup>-6</sup> |

**Table S3.** Genes at loci that influence growth in *mrp20-105E* segregants. These loci additively affect the expressivity of *mrp20-105E* mutation. Recombination delimits each peak to between one candidate (12 loci), two candidate genes (3 loci) or 4 genes (1 locus). Location of the delimited SNPs is included.

| Chromosome | Peak Position(s) | Candidate Gene(s) | Position of Peak SNP(s) |
| --- | --- | --- | --- |
| 4 | 615,114 | AFR1 | Coding |
| 7 | 140,806 to 142,398 | HOS2, YGL193C, and IME4 | Promoter and Coding |
| 11 | 171,261 to 171,457 | SDH1 | Promoter |
| 12 | 116,314 116,980 | BPT1 | Promoter and Coding |
| 12 | 448,368 | RNH203 | Promoter |
| 12 | 679,600 | BOP2 | Coding |
| 12 | 1,022,895 to 1,022,933 | ECM7 | Coding |
| 13 | 449,844 to 451,590 | YMR090W and NPL6 | Promoter |
| 13 | 612,355 | ECM5 | Coding |
| 13 | 833,534 | AEP2 | Coding |
| 14 | 299,515 | PBR1 | Coding |
| 14 | 472,584 to 473,648 | SAL1 and PMS1 | Promoter and Coding |
| 15 | 185,793 | BRX1 | 3' UTR |
| 15 | 343,484 to 343,921 | YOR008C-A and TIR4 | Promoter |
| 15 | 627,315 to 628,209 | ISN1 | Promoter and Coding |
| 15 | 885,437 to 885,914 | ISW2 | Promoter and Coding |

**Table S4.**
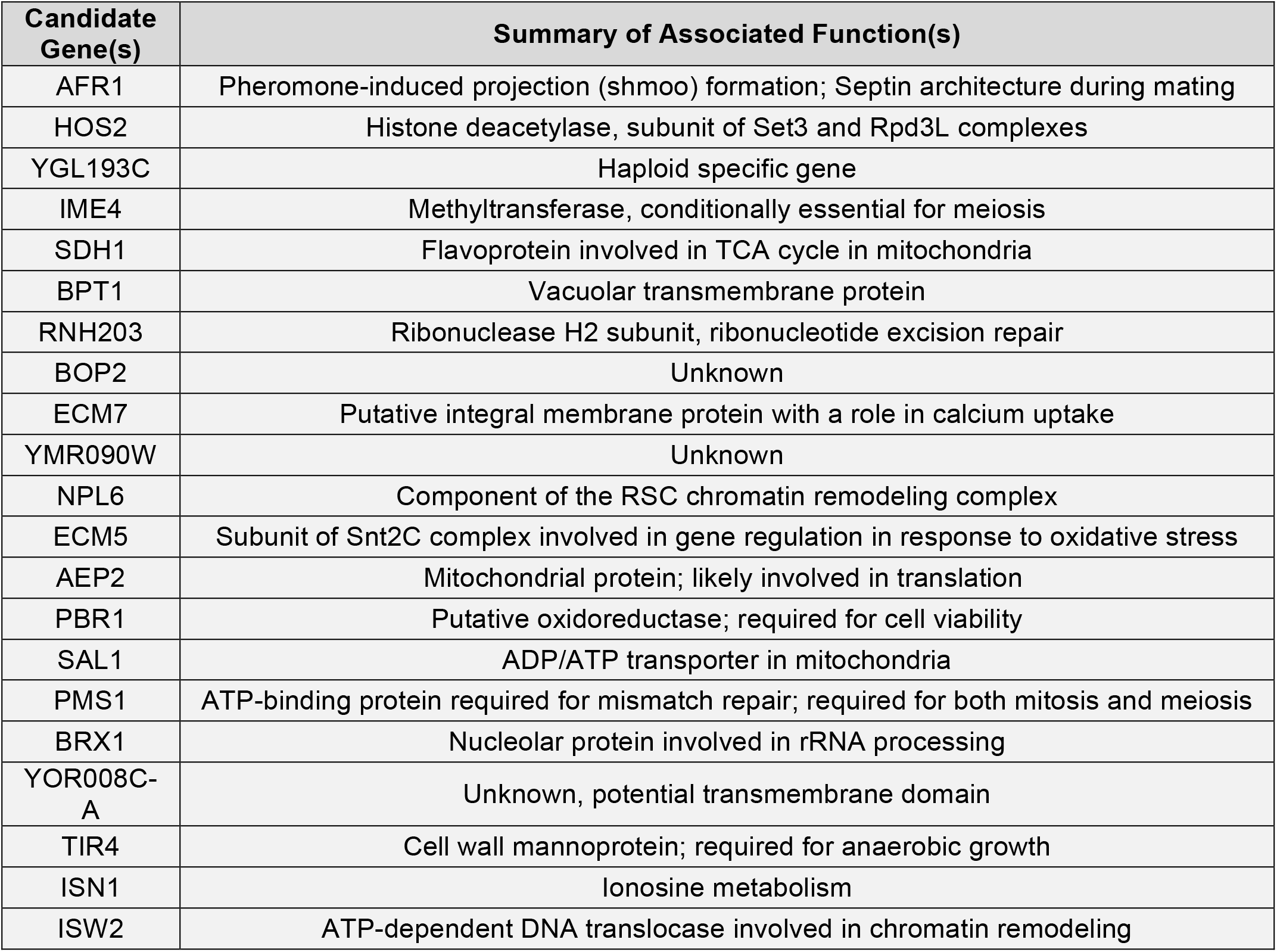
Candidate genes have diverse cellular functions. Candidate genes delimited by recombination at *mrp20-105E* listed in table S3 are included here with a brief summary of each’s function based on summarized descriptions on the *Saccharomyces* Genome Database.

| Candidate Gene(s) | Summary of Associated Function(s) |
| --- | --- |
| AFR1 | Pheromone-induced projection (shmoo) formation; Septin architecture during mating |
| HOS2 | Histone deacetylase, subunit of Set3 and Rpd3L complexes |
| YGL193C | Haploid specific gene |
| IME4 | Methyltransferase, conditionally essential for meiosis |
| SDH1 | Flavoprotein involved in TCA cycle in mitochondria |
| BPT1 | Vacuolar transmembrane protein |
| RNH203 | Ribonuclease H2 subunit, ribonucleotide excision repair |
| BOP2 | Unknown |
| ECM7 | Putative integral membrane protein with a role in calcium uptake |
| YMR090W | Unknown |
| NPL6 | Component of the RSC chromatin remodeling complex |
| ECM5 | Subunit of Snt2C complex involved in gene regulation in response to oxidative stress |
| AEP2 | Mitochondrial protein; likely involved in translation |
| PBR1 | Putative oxidoreductase; required for cell viability |
| SAL1 | ADP/ATP transporter in mitochondria |
| PMS1 | ATP-binding protein required for mismatch repair; required for both mitosis and meiosis |
| BRX1 | Nucleolar protein involved in rRNA processing |
| YOR008C-A | Unknown, potential transmembrane domain |
| TIR4 | Cell wall mannoprotein; required for anaerobic growth |
| ISN1 | Ionosine metabolism |
| ISW2 | ATP-dependent DNA translocase involved in chromatin remodeling |

**Table S5.** Presence of Chromosome II duplication differs among BY x 3S crosses. We observed a Chromosome II duplication in crosses fixed for *mrp20-105E*. Among these two crosses, the cross fixed for *MKT1^3S^* had much higher prevalence of the aneuploidy relative to the cross fixed for *MKT1^BY^*.

| Diploid | Cross | % wildtype | % aneuploid |
| --- | --- | --- | --- |
| A | BY x 3S | 100 | 0 |
| D | BY *mrp20-105E* *MKT1*<sup>B</sup> x 3S *mrp20-105E* *MKT1*<sup>BY</sup> | 94 | 5.9 |
| E | BY *mrp20-105E* *MKT1*<sup>3S</sup> x 3S *mrp20-105E* *MKT1*<sup>3S</sup> | 51 | 49 |

## Supplementary Data

**Dataset S1. Segregant genotype table**

Chromosome and position columns refer to the chromosome and position of each genetic variant used in this study, The mitochondria is referred to as chromosome 17. Each additional column contains the genetic information for a given segregant. Each segregant was named by type (F_2_ and F_3_), the diploid from which it originated (A, B, C, D, E defined in table S2), whether that segregant was wildtype or mutant at *MRP20* (‘MRP20’ or ‘mrp20’), and was randomly numbered from one through the total number of that segregant type. Segregants originating from diploid B contained additional information pertaining to whether that segregant was wildtype or knockout at *HOS3* (‘HOS3’ or ‘hos3Δ’) and whether that segregant was obtained by random spore prep or tetrad dissection (‘random’ or ‘dissected’). Note, that for *hos3*Δ segregants obtained by random spore preparation of diploid B, the BY allele at *MRP20* contained the *mrp20-105E* mutation. Each genetic variant used in this study is presented as a row, whereby the haplotype information for each segregant is denoted as 0 for BY or 1 for 3S respectively. A value of ‘NA’ indicates a site that lacked coverage for which a haplotype was not called. In the BY x 3S crosses fixed for *mrp20-105E* (diploids D and E), a third heterozygous state (2) was used to denote heterozygosity for individuals with the chromosome II duplication event.

**Data S2. Segregant phenotype table**

Each segregant’s growth in ethanol is presented. A single growth measurement is reported, which is the mean value of three biological replicates of growth normalized to on plate BY controls.

**Data S3. Reciprocal hemizygosity experiments**

Segregants that were used in reciprocal hemiygosity experiments that delimied the Chromosome IV allele to *MRP20* are included. The segregants mated together for each hemizygous diploid are listed under ‘Parent1’ and ‘Parent2’ columns. The ‘Gene’ column lists the gene at which reciprocal hemizygosity was engineered. The ‘LossOfFunction’ column indicates which allele, BY or 3S (encoded as 0 or 1) was engineered to be non-functional. The ‘Ethanol’ columns contains a growth value normalized to on plate BY controls. Each biological replicate is included separately and denoted by the ‘Replicate’ column to enable calculation of confidence intervals.

**Data S4. Cloning experiments**

Each segregant and parent strain used for cloning causal nucleotides at *MRP20* and *MKT1* are included. The ‘Type’ column denotes whether the engineered strain is a segregant or parent (‘segregant’ or ‘parent’), the ‘MRP20’ column describes whether that strain is wildtype or *mrp20-105E* (denoted as ‘MRP20’ or ‘mrp20’), and the ‘MKT1’ column describes whether that strain was BY or 3S (encoded as 0 or 1) at the causal SNP at position 467,219. The ‘Gene’ column lists the gene at which engineering had occurred, including (‘MRP20’, ‘MKT1’, ‘MRP20andMKT1’, and ‘WT’ which denotes parental control samples. The ‘Edit’ column explains the type of engineering that was performed in segregants. Thus, cloning experiments at *MRP20* in segregants are described as ‘fromMuttoWT’ or ‘fromWTtoMut’. Similarly cloning experiments at *MKT1* in segregants are described as ‘from3StoBY-1SNP’, ‘from3StoBYcandidate’, and ‘from3StoBY+1SNP’ for strain engineering at the nearest upstream, causal, and nearest downstream SNPs. This column is not relevant for parental cloning and therefore NA is reported in those cells. Lastly, the ‘Ethanol’ column contains a single growth measurement which is the mean value of three biological replicates of growth normalized to on plate BY controls.

**Data S5. Petite frequency**

Each strain used in petite frequency assays, either segregants described in Data S1-S3 or parental strains is reported in the ‘Sample’ column. The ‘Type’ column denotes whether the given strain is a segregant or parent (‘segregant’ or ‘parent’), and the ‘MRP20’ column describes whether or not the strain is wildtype or *mrp20-105E* (denoted as ‘MRP20’ or ‘mrp20’). A comma separated list of Image J reported colony sizes (in^2^) is also included in the ‘ColonySizes’ column. The largest observed petite colony among wildtype parental strains was 0.001 in^2^ and was thus used to designate *petites* from total colonies. The final ‘Frequency’ column reports the petite frequency, or the number colonies at or below this threshold relative to the total number of colonies times 100.

**Data S6.Individuals with Chromsome II duplication event**

Each segregant from BY x 3S crosses that were engineered at *mrp20-105E* and *MKT1* (croses D and E) in which the Chromosome II duplication was observed is listed under the ‘Sample’ column, The average normalized coverage acrpss chromosome II is reported in the ‘AverageChr2Coverage’ column, and the ‘Chr2C’ column denotes if that segregant was determined to be WT or Aneuploid (encoded as 0 or 1).

**Data S7. Genetic mapping analysis code**

All code used for linkage mapping and multiple testing correction is included as ‘.R’ file to be used in R programming language.

**Data S8. Statistical analysis code**

All code used for statistical analyses is included as ‘.R’ file used to be used in R programming language.

**Data S9. Figure plotting code**

All code used for plotting data shown in main text and manuscript is included as ‘.R’ file to be used in R programming language.

### Links to Supplementary Data

**DataS1:** https://drive.google.com/file/d/1OURXNu8bHkbB6rbUzswF07vBauYfmQ9S/view?usp=sharing

**DataS2:** https://drive.google.com/file/d/1_6LaPckKvzLdkSaeefjjQc7yAQ1L-NA6/view?usp=sharing

**DataS3:** https://drive.google.com/file/d/1TEmdHQ_5yHsrsiH-1qJZxSr4o0WSwVzd/view?usp=sharing

**DataS4:** https://drive.google.com/file/d/1k1-XgWtn2GPRsaPBTnrrGRwu9T4Z_jfS/view?usp=sharing

**DataS5:** https://drive.google.com/file/d/1u71y1TV7vdCjrl2dN72uAO1JUlXJunzq/view?usp=sharing

**DataS6:** https://drive.google.com/file/d/1OZNNJxVE5lLF3JgwdD5LlfE2TYbCPX7T/view?usp=sharing

**DataS7:** https://drive.google.com/file/d/1gUaZgfqJk4jq7AUliAH30VYYzcC42K8n/view?usp=sharing

**DataS8:** https://drive.google.com/file/d/1Wnvf-CurH8UceBTYJcE4S78IeWoHwUW6/view?usp=sharing

**DataS9:** https://drive.google.com/file/d/1emHITn8rnEB6Y8HpBp9kIG3nIHuMxwxE/view?usp=sharing

